# Acquired mutations and transcriptional remodeling in long-term estrogen-deprived locoregional breast cancer recurrences

**DOI:** 10.1101/2020.06.08.140707

**Authors:** Nolan Priedigkeit, Kai Ding, William Horne, Jay K. Kolls, Tian Du, Peter C. Lucas, Jens-Uwe Blohmer, Carsten Denkert, Anna Machleidt, Barbara Ingold-Heppner, Steffi Oesterreich, Adrian V. Lee

## Abstract

Endocrine therapy resistance is a hallmark of advanced estrogen receptor (ER) positive breast cancer. In this study, we performed DNA/RNA hybrid-capture sequencing on 12 locoregional recurrences after long-term estrogen-deprivation along with each tumor’s matched primary. Despite being up to 7 years removed from the primary lesion, most recurrences harbored similar intrinsic transcriptional and copy number profiles. Only two genes, *AKAP9* and *KMT2C*, were found to have single nucleotide variant (SNV) enrichments in more than one recurrence. Enriched mutations in single cases included SNVs within transcriptional regulators such as *ARID1A, TP53, FOXO1, BRD1, NCOA1* and *NCOR2* with one local recurrence gaining three *PIK3CA* mutations. In contrast to DNA-level changes, we discovered recurrent outlier mRNA expression alterations were common—including outlier gains in *TP63* (n=5 cases [42%]), *NTRK3* [n=5 [42%]), *NTRK2* (n=4 [33%]), *PAX3* (n=4 [33%]), *FGFR4* (n=3 [25%]) and *TERT* (n=3 [25%]). Recurrent losses involved *ESR1* (n=5 [42%]), *RELN* (n=5 [42%]), *SFRP4* (n=4 [33%]) and *FOSB* (n=4 [33%]). *ESR1-*depleted recurrences harbored shared transcriptional remodeling events including upregulation of *PROM1* and other basal cancer markers. Taken together, this study defines acquired genomic changes in long-term, estrogen-deprived disease, highlights longitudinal RNA-profiling and identifies a common endocrine-resistant breast cancer subtype with basal-like transcriptional reprogramming.

## INTRODUCTION

Hormone receptor positive breast cancer has served as a prototype for targeted therapy due to the well-established efficacy of estrogen deprivation. Largely because of these approaches, breast cancers are somewhat unique in that recurrences can occur years, sometimes decades following the primary diagnosis^1–4^. Given that the majority of patients receive long-term maintenance regimens of either a selective estrogen receptor modulator (SERM) or aromatase inhibitor (AI), recurrent breast cancers are often classified as estrogen-independent given their ability to thrive in an estrogen-deprived environment. Identifying the biological mediators that allow breast cancer cells to bypass their dependence on estrogen is a crucial step in understanding advanced breast cancer biology and defining novel therapeutic targets.

Defining these molecular processes in patient samples, however, has been challenging because of the logistics in obtaining well-characterized, longitudinally collected biospecimens. Nevertheless, shared features of more advanced breast cancers have emerged, such as relapsed tumors losing expression of ER and over 20% of metastatic ER-positive breast cancers acquiring mutations in *ESR1* that confer ligand-independent signaling^5–7^. Other largely accepted mechanisms of estrogen-independence are bypass activations of mitogenic pathways such as MAPK and PI3K through initiating FGFR, EGFR and IGF signaling and exploitation of the Rb-CDK-E2F axis^8–12^. Less well validated, more recently discovered mechanisms include *ESR1* fusions and amplifications^13,14^.

Recent studies analyzing multiple, longitudinally collected, pre and post-treatment samples have shown clonal evolution and selection in the context of targeted therapies^15–18^. Similar work analyzing hormone-receptor positive breast cancers have mainly been restricted to short-term pre/post neoadjuvant therapy analyses^19–22^. One of the most comprehensive genomic studies of this type was a multi-platform effort that characterized the clonal architecture of tumors after four months of AI therapy^23^. Although drastic clonal remodeling was observed at the DNA-level, few recurrent resistance mechanisms were appreciated. A more recent, large-scale study showed activating *ERBB2* mutations, MAPK activation and NF1 loss as mechanisms possibly driving endocrine resistance—with some of these alterations being confirmed in subsequent studies^24–27^. The majority of this work has notably been performed on metastatic tissues—whether or not some of these changes occur locally as a result of estrogen-independence before distant spread is unknown.

Thus, to better define both DNA and transcriptional changes that occur in long-term estrogen-independent tumors, we undertook a targeted analysis of DNA/RNA alterations in ∼1,400 cancer genes in 12 paired primary and locoregional recurrences from patients with ER-positive breast cancers that were documented as being treated with estrogen-depleting therapy. The median time to recurrence was 3.7 years, with the longest time to recurrence being over 7 years.

## RESULTS

### Expression and copy number changes in local recurrences

Dual hybrid-capture DNA/RNA sequencing was performed for 1,400 cancer genes on 12 paired primary and local recurrences from patients with ER-positive breast cancers that underwent continuous endocrine therapy (Table 1). RNA-seq data (Supplementary Table 1) underwent unsupervised hierarchical clustering of normalized RNA expression values which showed most patient matched pairs clustered transcriptionally with their matched primary—regardless of the length of disease-free survival (Figure 1A). Unlike a previous transcriptome-wide analysis of primary breast cancers and matched bone metastases^28^, there was no significant correlation in pair transcriptional similarity and time to recurrence—although a trend towards negative correlation was observed (pearson R = −0.37, p-value = 0.236). Only a single recurrence showed marked transcriptional deviation from its matched primary (ERLR_03_R1); whereby it lost ER-positivity and gained HER2-positivity clinically. Copy number alterations (CNAs) between primary and recurrences were analyzed in the targeted capture regions for 10 cases (Supplementary Figure 1 and Supplementary Figure 2). Similar to expression, CNAs were largely consistent among the recurrences when compared to their matched primary (Figure 1B). Two exceptions were recurrences from cases ERLR_01 and ERLR_03, which showed distinct copy number profiles from the matched primary tumors with poor correlation between primary and recurrence CNA values versus all other cases (Supplementary Figure 3). Notably, unlike case ERLR_03, ERLR_01 interestingly retained a similar expression profile despite a distinct CNA profile. An analysis of shared variants validated both DNA and RNA extracts originated from the same patient (Supplementary Figure 4), excluding the possibility of sample mixup. *ERBB2* copy number values correlated well with RNA expression (pearson R 0.92), as did other amplified genes including *CDK12* and *CCND1* (Figure 1C, Data Supplement S3).

**Table 1:**
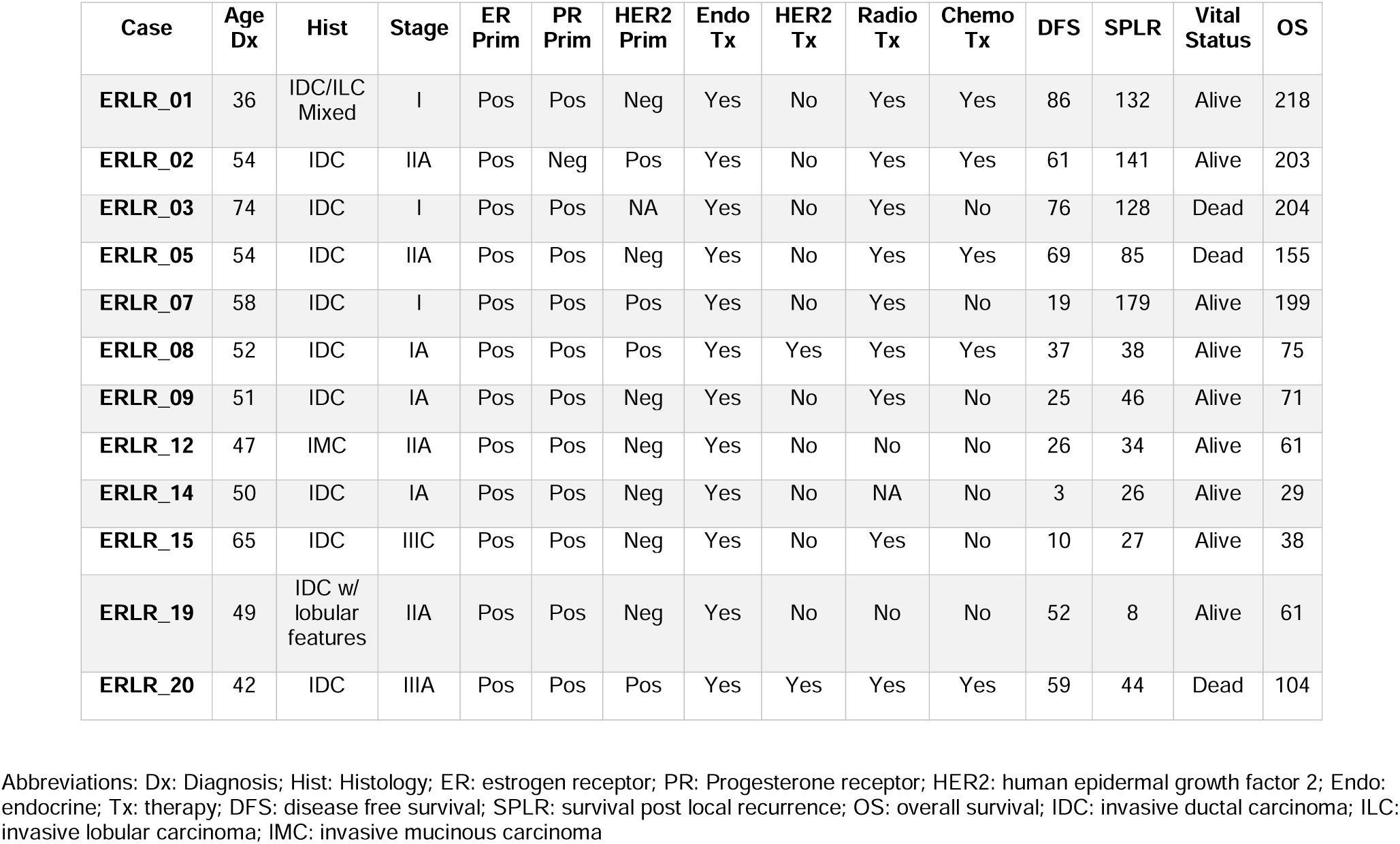
Abridged clinicopathological features of patient-matched primary and local recurrence tumor cohort^¥^.

**Figure 1:**
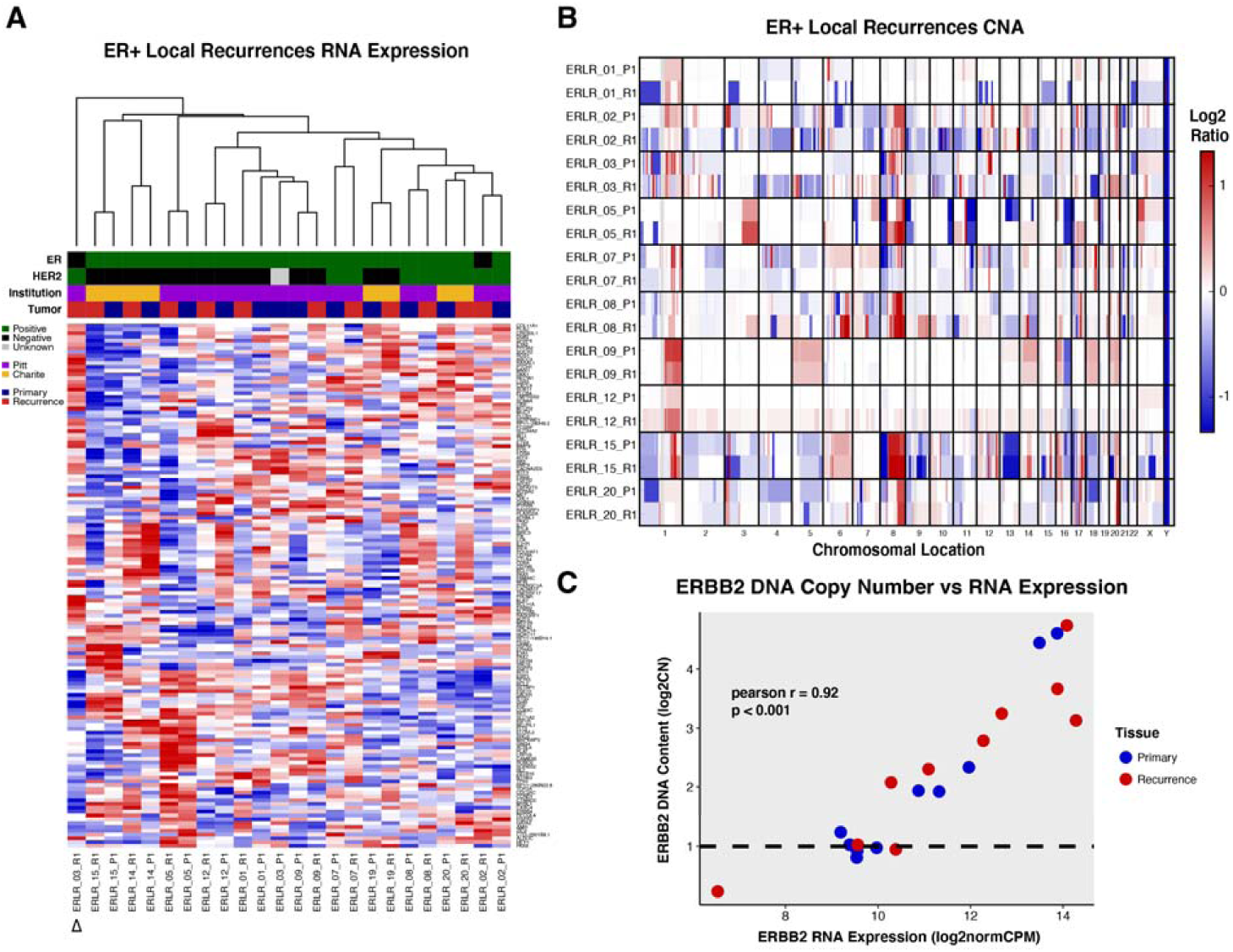
Transcriptional and CNA profiles of ER-positive local recurrences. **(A)** Unsupervised hierarchical clustering and heatmap (red = high relative expression, blue = low relative expression) on normalized gene expression values from patient-matched pairs (P1 = Primary, R1 = Recurrence). Clinical ER and HER2 status (black = negative, green = positive, grey = unknown), tissue source site (purple = Pitt, yellow = Charite), and tumor type (blue = primary, red = recurrence) are indicated. Delta symbol shows distinct clustering of ERLR_03_R1 away from its matched primary, ERLR_03_P1. **(B)** Heatmap of copy number ratios from patient-matched pairs. Redder regions indicate regions of copy number gain and bluer regions indicate regions of loss. **(C)** Correlation between ERBB2 DNA copy number calls and normalized expression values.

### SNV enrichments and differentially expressed genes

A total of 406 distinct, presumed-somatic nonsynonymous mutations were detected in either a primary or recurrence at an AF > 5% among the 10 DNA-sequenced cases (Data Supplement S4). To assess if there are shared DNA mutations acquired in recurrences, an analysis of enriched single nucleotide variants (SNVs) was performed which showed 56 statistically enriched SNVs in local recurrences versus matched primary tumors (Figure 2A, Data Supplement S5). SNVs in two genes were found to be enriched in more than one case (n = 2 [20%]), *AKAP9* (R3320W, S319*) and *KMT2C* (T1969I, Y366N, R894Q). The recurrent mutations did not exhibit features suggesting functional selection, such as being within a conserved functional domain or within a COSMIC^29^ hotspot region, making it difficult to assess if these are pathogenic. Other case-specific, n-of-one enriched mutations included nonsense mutations in *ARID1A* (Q1424*, Case ERLR_20, Primary AF 0.5%, Recurrence AF 16.5%) and *BRD1* (Q467*, Case ERLR_01, Primary AF 0.93, Recurrence AF 57.88%), an acquired *TP53* mutation (S241C, Case ERLR_03, Primary AF 0.0%, Recurrence AF 53.4%) and an enriched *NCOR2* mutation (A4942C Case ERLR_08, Primary AF 4.4%, Recurrence AF 19.4%). In case ERLR_01, an enrichment of a suite of three somatic mutations in *PIK3CA* was observed (E542K, Q546K, E726K) in the recurrence (Figure 2B). Notably, the number of enriched, non-silent SNVs ranged from 0 to 13 and was positively correlated with clinical time to recurrence (Figure 2C). No acquired *ESR1* mutations were observed. These mutations were examined in the corresponding RNA-seq data to determine if they are expressed. Out of 633 total mutations—considering some of the 406 distinct mutations were present in both matched tumors—315 were detected in RNA with at least 2 supporting reads of the altered allele and an AF >=5%. Allele frequencies called from DNA-seq and RNA-seq data correlated well (Supplementary Figure 5, pearson R = 0.609, p-value=2.57e-33). Noteworthy, out of the 56 enriched SNVs in recurrence at DNA level, 31 distinct mutations can be detected with confidence at the RNA level (Data Supplement S6)—including *AKAP9, KMT2C, ARID1A, BRD1* and *TP53* as discussed above.

**Figure 2:**
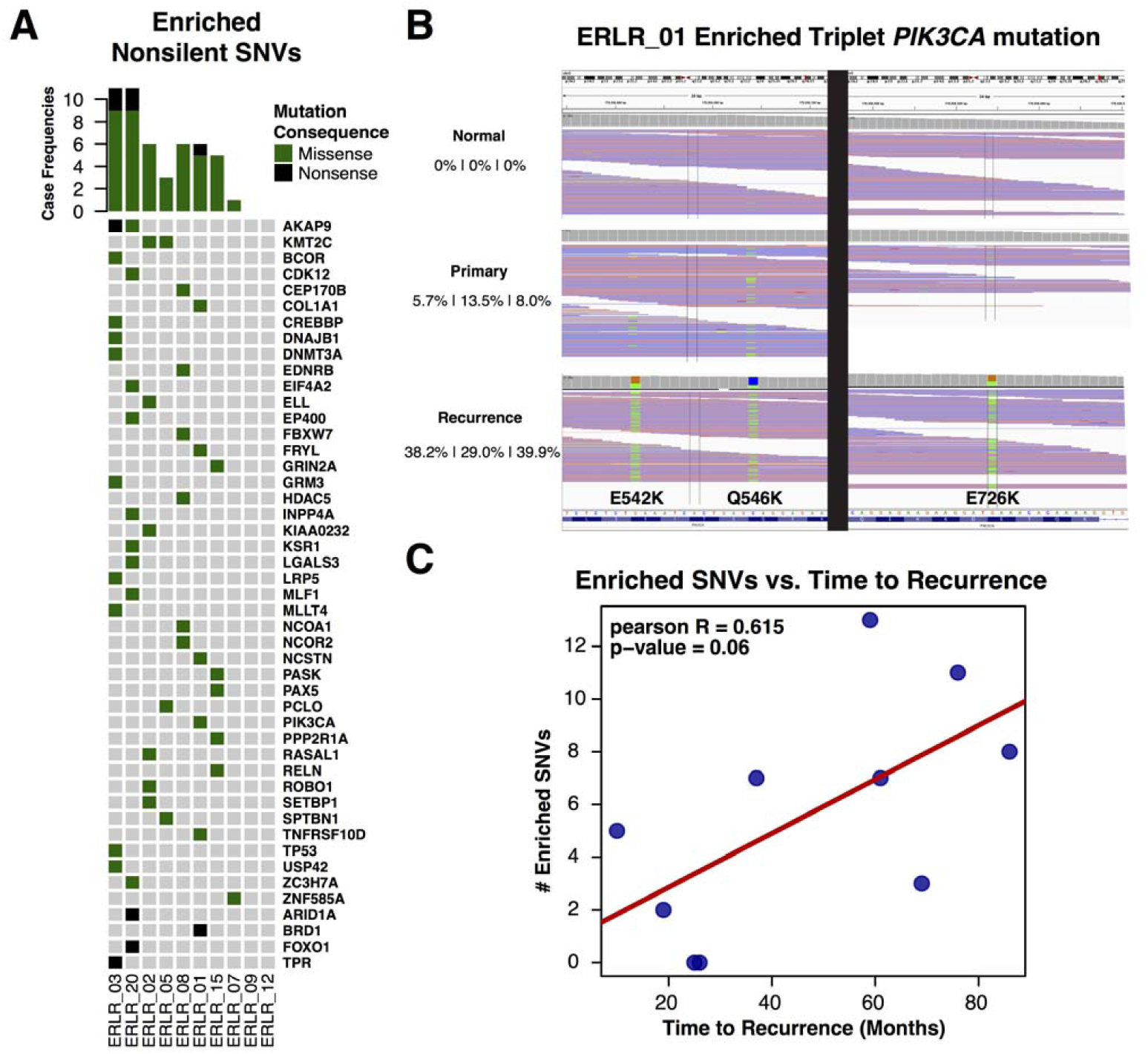
SNV enrichments in ER-positive local recurrences. **(A)** OncoPrint of non-silent, enriched single nucleotide variants in patient-matched cases. Missense variants are indicated with a green box and nonsense variants with black. **(B)** Triplet mutation enrichment of PIK3CA mutations in case ERLR_01. Collapsed IGV alignments are shown, along with allele frequencies, for the normal, primary and recurrence. **(C)** Frequency of enriched, non-silent single nucleotide variants versus time to recurrence along with pearson R and calculated p-value.

A differential expression analysis, comparing all primary tumors versus all local recurrences, yielded no genes passing an FDR corrected p-value of less than 0.05—which is perhaps expected given heterogeneity of clinical specimens (Data Supplement S7). Nonetheless, 71 genes with an average, *voom* normalized expression value of 2 or greater, a nominal p-value of less than 0.05 and a log2 fold-change greater than +/- 0.5 were identified (Table 2). Some of these genes, including the upregulation of *EPOR, NDRG1, IDH2, CEBPA* and *PTPA* and downregulation of *ESR1, IGF1R, NFKB1* and *RUNX2*, are also differentially expressed in long-term estrogen deprived ER-positive cell lines (Supplementary Figure 6)^30^.

**Table 2:** Differentially expressed genes in ESR1 depleted recurrences.

| Gene | Log2FC | voom Average Expression | Nominal P-value | FDR Adjusted P-value |
| --- | --- | --- | --- | --- |
| *PAPPA* | 1.395 | 6.416 | 0.001 | 0.293 |
| *KLK7* | 5.422 | 1.158 | 0.001 | 0.293 |
| *PROM1* | 3.931 | 5.005 | 0.002 | 0.588 |
| *RASGRF1* | 2.307 | 4.106 | 0.002 | 0.588 |
| *DKK1* | 2.732 | 0.473 | 0.004 | 0.614 |
| *EPHB6* | 1.641 | 3.819 | 0.005 | 0.614 |
| *ABCC3* | 1.637 | 8.010 | 0.006 | 0.614 |
| *FGFR4* | 1.515 | 5.267 | 0.010 | 0.695 |
| *FBN2* | 1.010 | 5.326 | 0.010 | 0.695 |
| *TENM1* | 1.326 | 4.709 | 0.012 | 0.705 |
| *COL9A3* | 2.034 | 2.249 | 0.014 | 0.705 |
| *NDRG1* | 1.218 | 8.945 | 0.014 | 0.705 |
| *TP63* | 2.135 | 4.441 | 0.018 | 0.768 |
| *SCN8A* | 1.290 | 5.881 | 0.019 | 0.768 |
| *KIT* | 1.289 | 6.020 | 0.020 | 0.768 |
| *TCL6* | 2.228 | -0.254 | 0.022 | 0.790 |
| *WNT11* | 1.585 | 1.256 | 0.024 | 0.823 |
| *SOCS1* | 1.534 | 0.387 | 0.033 | 0.911 |
| *HOXD11* | 2.755 | -1.369 | 0.034 | 0.911 |
| *PLAG1* | 1.275 | 4.576 | 0.036 | 0.911 |
| *DTX4* | 1.185 | 5.711 | 0.036 | 0.911 |
| *FLNC* | 1.588 | 6.787 | 0.037 | 0.911 |
| *ALDOC* | 1.494 | 5.224 | 0.039 | 0.911 |
| *ACSBG1* | 1.843 | 0.601 | 0.042 | 0.915 |
| *SYP* | 1.348 | 0.862 | 0.045 | 0.915 |
| *ESR1* | -3.952 | 9.492 | 0.000 | 0.146 |
| *ATP8A2* | -2.599 | 4.510 | 0.003 | 0.588 |
| *ELOVL2* | -2.090 | 2.413 | 0.006 | 0.614 |
| *RABEP1* | -1.009 | 10.352 | 0.012 | 0.705 |
| *EYA1* | -1.494 | 2.203 | 0.013 | 0.705 |
| *IGF1R* | -1.149 | 9.083 | 0.016 | 0.747 |
| *CAMK2A* | -1.391 | 2.742 | 0.016 | 0.747 |
| *RERG* | -1.413 | 6.562 | 0.018 | 0.768 |
| *BCL2* | -1.055 | 6.619 | 0.020 | 0.768 |
| *FGF14* | -1.393 | 2.430 | 0.023 | 0.790 |
| *RASGRP1* | -1.044 | 6.799 | 0.027 | 0.857 |
| *BHLHE22* | -1.822 | 0.822 | 0.035 | 0.911 |
| *ZNF703* | -1.811 | 4.865 | 0.038 | 0.911 |
| *MYB* | -1.179 | 8.857 | 0.045 | 0.915 |

### Outlier expression gains and losses

To further explore major expression changes that may be driving recurrence and therapy resistance, an outlier expression analysis was performed using gene-level fold-change values of each patient-matched case (Data Supplement S8). Unlike non-silent SNVs, recurrent transcriptional gains and losses were common (Figure 3A). These included gains and losses in shared pathway members, notably *NTRKs* and *SFRPs* respectively, targetable upregulation of growth factor pathway mediators such as *FGFR4* and *EGF* and outlier gains in the CDK regulator *CCNE1*. 3 of 12 cases also shared outlier expression gains in *TERT*, with case ERLR_14 harboring a particularly extreme enrichment from near undetectable levels in the primary tumor (Figure 3B). Case ERLR_03’s recurrence, which was most dissimilar to its patient-matched pair transcriptionally, showed extreme loss and gain of *ESR1* and *ERBB2* respectively. CNA analysis confirmed recurrence-specific *ERBB2* amplification and is consistent with previous studies of endocrine therapy-treated breast cancers selecting for HER2-signaling in more advanced tumors. The most recurrent outlier loss involved *ESR1*.

**Figure 3:**
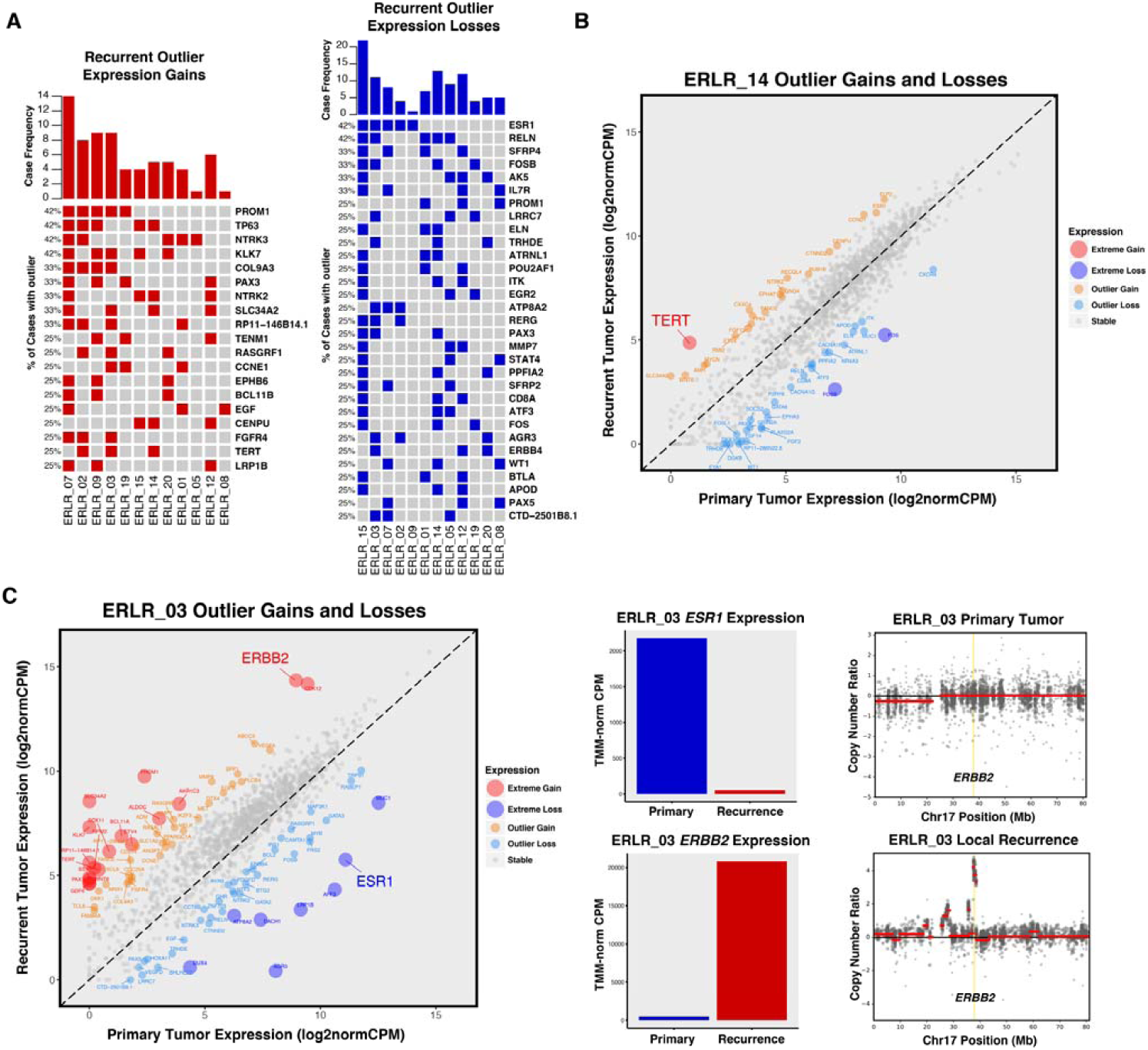
Outlier expression gains and losses in ER-positive local recurrences. **(A)** OncoPrint of outlier expression gains (red) and outlier expression losses (blue) in ER-positive local recurrences. Genes are sorted by frequency of outlier changes across pairs. **(B)** Extreme expression gain of TERT in case ERLR_14; 2 other cases showed similar TERT enrichments in recurrent tumors. **(C)** Extreme expression gain and loss of ERBB2 and ESR1 respectively. TMM-normalized CPM values of primary (blue) and recurrent (red) tumor. ERBB2 expression gain is driven by recurrence-specific DNA-level amplification of ERBB2 locus.

### ESR1 depleted recurrences

Five cases showed outlier expression losses of *ESR1* (Figure 4A). Despite estrogen receptor being the driver of ER-positive breast cancer and a major regulator of transcription; counterintuitively, 4 of 5 of the recurrences which lost *ESR1* expression generally retained the expression profile of their patient-matched primary (Figure 1A). Importantly, many of these cases also harbored very similar CNA profiles (Figure 1B), implying the recurrences were derived from a continuous clonal lineage as opposed to being completely distinct breast cancers. Thus, to explore the transcriptional consequences of acquired *ESR1* loss in ER-positive disease and identify potential bypass mechanisms driving ER*-* independence, a differential expression analysis was performed on the subset of pairs with outlier *ESR1* expression losses. This analysis revealed several recurrently dysregulated genes in *ESR1* depleted recurrences (Data Supplement S9). Two standout genes, *KLK7* and *PROM1*, showed the highest degree of fold change with a log2 fold-change increase of 5.4 and 3.9 respectively—with some tumors exhibiting changes from near undetectable levels to high expression (Figure 4C). These two genes are more commonly expressed in basal cancers, with *PROM1* being a cancer lineage stem cell marker (Supplementary Figure 7)^31^. Other genes with significant log2 fold-changes > 1 included drug targets such as *FGFR4, KIT, IGF1R* and *BCL-2* (Table 2). *NDRG1*, a particularly compelling candidate since it also showed upregulation in LTED breast cancer models, was further interrogated using METABRIC data. Like *PROM1* and *KLK7, NDRG1* is most highly expressed in basal breast cancers; yet, when expressed in ER-positive primary tumors, *NDGR1* confers significantly worse disease-specific survival outcomes (Supplementary Figure 8).

**Figure 4:**
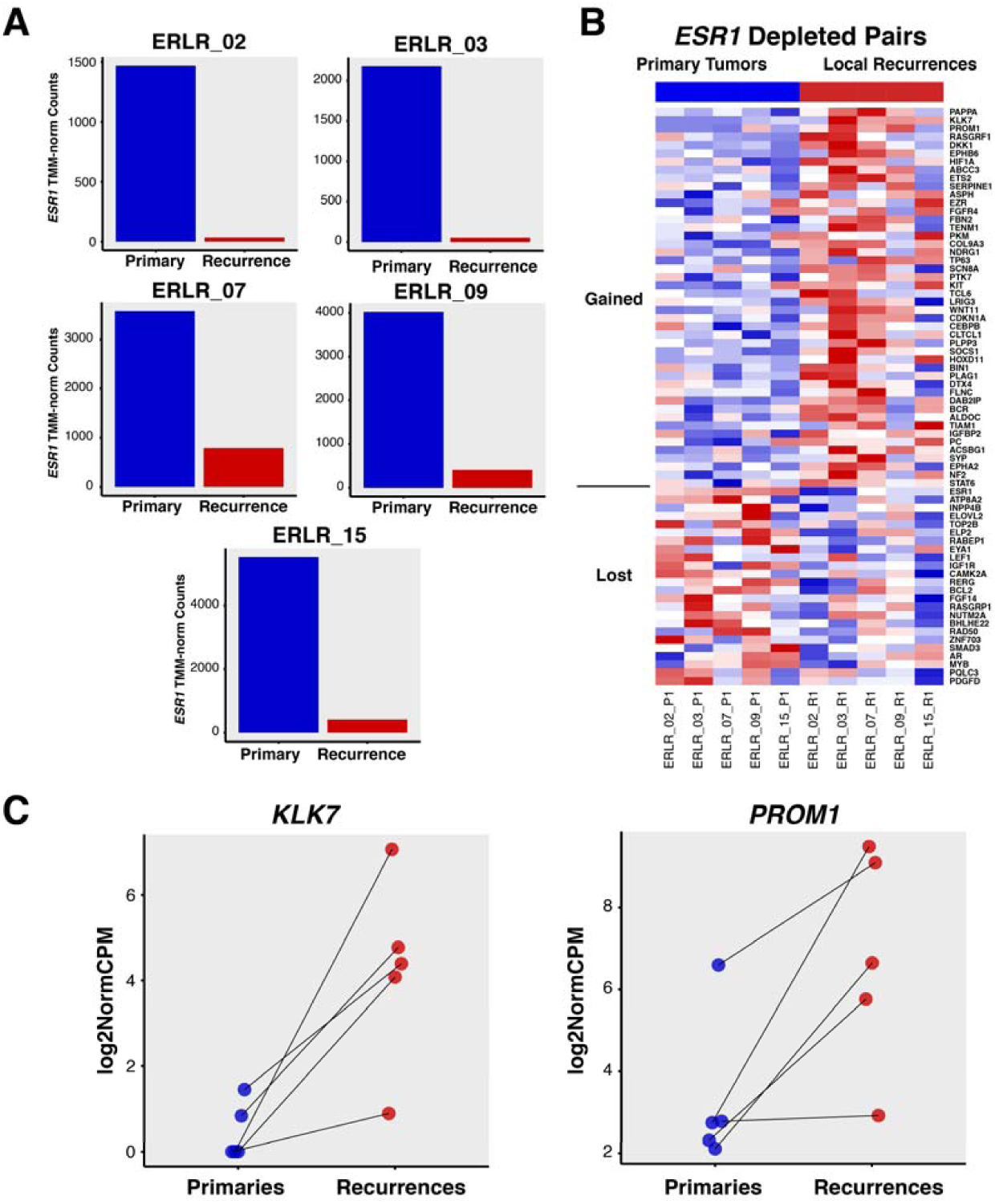
ESR1 depleted recurrences. **(A)** TMM-normalized expression of patient-matched local recurrences; primary tumor expression in blue, recurrent tumor expression in red. **(B)** Heatmap of differentially expressed genes (nominal p-value < 0.05, red = high relative expression, blue = low relative expression) in ESR1 depleted recurrences versus matched primary tumors. Genes are sorted by p-value and segregated by log2 fold-change values; log2 fold-change > 0 on top, log2 fold-change < 0 on bottom. (C) Ladder plots showing log2normCPM expression values for both KLK7 and PROM1, two of the most significantly upregulated genes in local recurrences with the largest average log2 fold-changes.

## DISCUSSION

In this study, a targeted RNA/DNA analysis of approximately 1,400 cancer genes in ER-positive primary breast cancers and matched long-term, endocrine therapy treated local recurrences was performed. We found general conservation of transcriptional and copy number profiles among the majority of samples—suggesting that even after 7 years of dormancy and the selective pressures of therapies, locally recurrent breast cancers generally retain their intrinsic molecular features. An analysis of recurrence-enriched SNVs revealed limited recurrent mutation events, yet notable “n-of-one” mutation selection was observed—such as case ERLR_01 which showed three distinct, recurrence-enriched *PIK3CA* mutations. The most striking changes in long-term estrogen-deprived tumors; however, were highly recurrent (up to 42%), outlier expression changes. An analysis of tumors with the most recurrent outlier loss, *ESR1*, revealed concurrent upregulation of genes typically expressed in basal breast cancers, such as *PROM1, KLK7* and *NDGR1*, suggesting selection of a more basal-like phenotype in endocrine-resistant disease. Our data showing similar CNA profiles argue against the outgrowth of a distinct ER-negative subclone but instead suggest possible epigenetic, transcriptionally-driven remodeling under antiestrogen pressures.

Nearly all recurrences are more similar transcriptionally to their matched primaries than to other, long-term estrogen deprived tumors—reinforcing the notion that advanced cancers generally retain their core transcriptional programming, even after nearly a decade of dormancy^26-29^. Furthermore, amplifications and deletions of recurrences are markedly similar to primaries, supporting recent evidence from breast cancer single-cell sequencing that structural variation is likely an early event and many CNAs, even in metachronous therapy-resistant tumors, may be shared by the majority of subclones^32^. An important exception to this conservation was ERLR_03_R1, a recurrence with a completely unique transcriptional and copy number profile than its matched primary. Evidence has emerged of so-called ‘collision tumors’, whereby two synchronous, distinct cancers can merge anatomically and only under the selective pressures of therapy or through deep sequencing, their individuality can be unmasked^23,33^. Indeed, this “recurrence” switched to ER-negative/HER2-positive from ER-positive/HER2-negative clinically, and thus could represent a different cancer than the primary— although the level of shared SNVs suggests some degree of clonal relatedness.

Limited shared, non-silent SNVs were discovered in these specimens, with *AKAP9* and *KMT2C* being the only two genes that harbored recurrence-enriched mutations in greater than one case. These mutations are not in a conserved functional domain nor in a hotspot location, making it difficult to assess their pathogenic roles. *AKAP9* and *KMT2C* also encode relatively large gene products (3911 and 4911 amino acids, respectively) which may increase the likelihood of obtaining a passenger mutation by chance. Nevertheless, *KMT2C* and other lysine methyltransferases have been implicated in breast cancer pathology, argued as potential drivers in large-scale sequencing studies of primary tumors and *KMT2C* mutations specifically may confer hormone therapy resistance in breast cancer models^34–36^. Case ERLR_20 harbored an enriched nonsense mutation in *ARID1A. ARID1A* alterations are associated with more unfavorable tumor features in breast cancer, and has recently been shown to determine luminal identify and therapy response in ER-positive tumors—consistent with the more basal-like transcriptional features we observe with ESR1-depleted recurrences^37–40^. A single recurrent cancer (ERLR_01_R1) showed enrichment of three somatic hotspot *PIK3CA* mutations (E542K, Q546K, E726K), suggesting strong MAPK signaling selection within that particular tumor and coincident with recent reports of multiple mutations occurring in individual cancer genes in advanced cancers^41^. SNVs within genes that act as corepressors and coactivators, some with direct influences on estrogen receptor mediated transcription, were found to be enriched in recurrences—such as *NCOA1, NCOR2, FRYL* and *CREBBP*—along with transcription factors including *PAX5, FOXO1* and *TP53*. Notably, we did not observe any *ESR1* mutations unlike other studies on locoregional recurrences^42^—likely due to our small sample size. Interestingly, this study reported lower frequency of *ESR1* mutations in locoregional recurrences versus advanced metastases at an AF > 1% and recent data has emerged regarding a pro-metastatic phenotype of *ESR1* variants^43^—suggesting locoregional recurrences may have a lower frequency of *ESR1* variants versus distant disease. We also observed a positive correlation between the frequency of acquired, non-silent SNVs and disease-free survival—validating the concept that surviving cancer cells after initial therapy acquire potentially pathogenic mutations as they lay dormant and undetectable over time.

Given the heterogeneity of clinical specimens makes it difficult to rely on typically used differential expression workflows—since resistant mechanisms of individual tumors may be distinct—we undertook an analysis of patient-specific outlier expression gains and losses to identify more extreme transcriptional reprogramming events within individual cases that may be driving estrogen independence. Surprisingly, unlike SNVs, recurrent outlier transcriptional gains and losses were quite common. Particularly compelling outlier events included recurrent gains within shared pathway members, such as near mutually exclusive upregulations of *NTRK3* [n = 5 [42%]) and *NTRK2* (n = 4 [33%]). Notably, activation of *NTRK’s* mediates downstream signaling pathways typically associated with breast carcinomas, including PI3K and MAPK, and small molecule inhibitors of this family are showing promising results in recent solid tumor trials^44^. Other notable pathway member changes included loss of Wnt antagonists *SFRP2* (n = 3 [25%]) and *SFRP4* (n = 4 [33%]). *SFRP2* is hypermethylated and silenced in a subset of breast cancers^45^ and experiments in model systems have shown cross-talk between ER and Wnt signaling that may mediate endocrine therapy resistance^46,47^. Other recurrent gains included *FGFR4* (n = 4 [33%]), *TERT* (n = 3 [25%]) and *CCNE1* (n = 3 [25%])*—*particularly relevant given the recent success of CDK inhibitors in hormone-positive disease and the burgeoning use of *FGFR* inhibitors against solid malignancies as we and others have reported^48,49^.

The most recurrent outlier expression loss was *ESR1*, which was diminished in 42% of long-term estrogen-deprived local recurrences. Interestingly, the loss of *ESR1* for the majority of cases was not associated with a dramatic change in the tumors’ transcriptional profile. To further explore this counterintuitive result, given *ESR1* is a master regulator of transcription and a driver of luminal breast cancers, we identified genes that were consistently altered in *ESR1* depleted recurrences. The most substantial gains in *ESR1* depleted tumors are genes generally expressed in basal breast cancers—such as *NDRG1, DKK1, KIT, KLK7, PROM1* and *COL9A3*—and genes significantly lost in the *ESR1* depleted subset are generally downregulated in basal cancers—*EVLOVL2, BCL2, IGF1R, MYB, RABEP* and *ATP8A2* (MsigDB: SMID_BREAST_CANCER_BASAL_DN/UP gene lists)^50^. These results reveal a common, novel and distinct *ESR1*-depleted subtype of advanced breast cancers that acquire basal-like transcriptional reprogramming.

The greatest fold-change difference in *ESR1* depleted recurrences was the upregulation of *PROM1. PROM1* is a marker for tumor-initiating cancer stem cells and plays a key role in determining ER-positive luminal cell fate during differentiation from multipotent stem cells^31^, suggesting long-term endocrine deprived breast cancer cells may enrich themselves with stem-like progenitors to achieve estrogen-independence. Indeed, *PROM1* has been shown to mediate endocrine therapy resistance in breast cancer models through IL6/Notch3 signaling ^51,52^. Here, we show that a large portion of long-term endocrine resistant breast cancers may be exploiting this transcriptional reprogramming. Finally, *NDRG1*, also significantly upregulated in *ESR1* depleted recurrences and generally expressed in basal cancers, showed differential expression in three distinct LTED cell lines. *NDRG1* is a suspected metastasis suppressor gene. Counterintuitively, we see upregulation of this gene in resistant disease and show increased expression confers worse survival outcomes in ER-positive primary tumors^53^. Further functional studies assessing the mechanistic and biological consequences of these transcriptional reprogramming events will be essential.

A pertinent point these results raise is the benefit of integrating longitudinal, targeted RNA-sequencing to inform resistance mechanisms and therapeutic targets in breast cancers. In this study, we find limited DNA-level enrichments yet highly recurrent, acquired transcriptional remodeling events from primary to advanced cancers, including a few of which that are immediately targetable such as *NTRKs, FGFR4* and *CCNE1.* Overall, this work challenges our lack of emphasis on RNA-level changes, particularly those that can be elucidated from longitudinal biopsies, in clinical profiling of tumors and future work should be geared towards deciphering which of these bypass transcriptional programs may be druggable.

Collectively, these results begin to unravel the complex adaptations that breast cancer populations undergo when under the selection of long-term estrogen depleting therapies long-term. We identify acquired DNA-level mechanisms of resistance, such as mutations in *ARID1A*, other transcriptional regulators and multiple mutation selection within *PIK3CA*—but more importantly, uncover the most recurrent genomic adaptations taking place appear to be at the transcriptional level. These include targetable outlier gains and modifications in *NTRKs* as well as a distinct population of *ESR1* depleted recurrences that enrich themselves with genes generally expressed in basal breast cancers—such as *PROM1* and *NDRG1*. Preclinical, mechanistic investigations into these temporally altered genes are warranted given they may uncover novel and targetable mechanisms of endocrine therapy resistance in advanced breast cancers.

## METHODS

### Patient Samples, tissue processing and nucleic acid extraction

Institutional Review Board approval from both participating institutions (University of Pittsburgh IRB# PRO15050502, The Charité IRB Office) was obtained prior to initiating the study. Inclusion criteria for this study were (1) patients harbored patient-matched formalin-fixed paraffin-embedded (FFPE) tissue from primary breast cancers and local recurrences (Table 4), (2) biospecimens contained macrodissectable regions with sufficient tumor cellularity and (3) disease was treated continuously with a form of estrogen-depleting therapy prior to the recurrence. Biospecimens were reviewed by a trained molecular pathologist to confirm pathology, quantify tumor cellularity and to highlight regions of relatively high tumor cellularity for macrodissection. If a slide region harbored sufficient, microscopically verifiable adjacent normal cells, this region was also dissected and processed for downstream analyses. Between four to ten (depending on tumor size) 10-micron FFPE sections immediately adjacent to the H&E-analyzed section were pooled and underwent dual DNA/RNA extraction using Qiagen’s AllPrep kit. Nucleic acids were quantified fluorometrically with a Qubit 2.0 Fluorometer and quality assessed with an Agilent 4200 TapeStation Instrument prior to sequencing.

### RNA and DNA-sequencing

RNA-seq library preparation was performed for all 12 cases using approximately 100 ng of RNA and Illumina’s *TruSight RNA Pan-Cancer* (1385 targets) protocol. DNA-seq library preparation was performed for 10 (6 with associated normal tissue) cases using no less than 30 ng of DNA and Illumina’s *TruSeq Exome* protocol with *TruSight RNA Pan-Cancer* probes for hybridization-based capture. Indexed, pooled libraries were then sequenced on Medium Output flow cells using an Illumina NextSeq 500 system (paired-end reads, 2 × 75 bp). A target of 5-10 million reads per sample was used to plan indexing and sequencing runs for RNA-sequencing and a target of 10-15 million reads was used for DNA-sequencing. RNA-sequencing FASTQ files were quantified with k-mer based lightweight-alignment (*Salmon* v0.7.2, quasi-mapping mode, 31-kmer index using GRCh38 Ensembl v82 transcript annotations, seqBias and gcBias corrections)^54^. *tumorMatch* (Chapter 3, Chapter 4) was used to validate sequencing pairs were patient-matched.

### RNA-sequencing and quantification and DNA-sequencing alignment

RNA-seq read counts and mapping percentages were calculated (Data Supplement 4: S1) and transcript abundance estimates were collapsed to gene-level with tximport^55^. Log2 transformed TMM-normalized CPM (log2normCPM) values were implemented for subsequent analyses^56,57^. DNA-seq reads were aligned with *bwa –mem* (v.0.7.13) to an hg19 reference, sorted with *samtools* (v1.3), duplicates marked and removed with *picardtools* (v1.140) and local realignment performed with *GATK* (v3.4-46)^58–60^. Average coverage depth for the processed bam file was calculated using *GATK’s DiagnoseTargets* and the Illumina *Pan-Can* bed file (Appendix A.4: Figure 41, Data Supplement 4: S2). Metrics for average coverage values across all target intervals were plotted with *ggplot2.*

### DNA-seq recurrence enriched variant determination

To determine enriched variants in recurrences versus patient-matched primary tumors, *VarScan2* was implemented^61^. More specifically, primary and recurrent *samtools* mpileup files derived from processed bam files were input into *VarScan2* using *somatic* mode, with somatic p-values representing the significance of a particular variant being acquired or enriched in the recurrence [SS = 1 or SS = 2]. Tumor purity estimates, as assessed by a molecular pathologist, were included in *VarScan2* to correct contaminating normal cell influence on allele frequencies. The minimum coverage for a variant to be considered was 40X, with a minimum allele frequency (AF) of 0.05 in either the primary or recurrence and a minimum of 5 reads supporting the variant. Germline variants were determined for cases containing a matched normal (ERLR_01, ERLR_02, ERLR_07, ERLR_08, ERLR_12 and ERLR_15) using *VarScan2*’s *germline* mode with the same parameters. VCF output files were then imported into R using the *VariantAnnotation* package^62^. If a normal sample was available for the case, all germline variants (AF > 0.30) were excluded from subsequent analyses. Additionally, to limit technical artifacts especially considering specimens were formalin-fixed paraffin embedded^63^, a “blacklist” of variants was created including those called in at least 3 of the normal samples. Germline and blacklist-removed variants were then annotated with *Annovar*^64^. Lastly, to call recurrence-enriched, potentially pathogenic variants the following inclusion criteria were enacted: (1) VarScan2 somatic p-value < 0.05, (2) > 2-fold gain in allele frequency in the recurrence versus the primary, (3) minimum AF of 0.10 in the recurrence, (4) non-silent and (5) an ExAC AF < 0.01 considering some samples were without a paired normal (Data Supplement 4: S3)^65^. These non-silent, enriched, potentially pathogenic variants were then plotted using the *OncoPrint* function in *ComplexHeatmaps*^66^. A pearson R correlation was calculated between the frequency of enriched variants and disease-free-survival. *PIK3CA* mutations were visualized with *IGV* (2.3.60)^67^ and variant allele frequencies were derived from *VarScan2*.

### RNA-seq variant determination

RNA-seq reads covering mutation sites called from DNA-seq of the corresponding sample were extracted from bam file and counted. Variants with at least 2 supporting reads containing the altered allele, and with AF greater than 0.05 in either primary or recurrence were considered.

### Copy number alterations

To estimate copy number ratios, *CNVkit* was implemented on processed bam files using default settings and the *-drop-low-coverage* option^68^. A pool of bam files from adjacent normal tissue, sequenced in the same manner, was used as a reference. Probe and segment level copy number estimates were finalized with *CNVkit*’s *call* function, which utilizes circular binary segmentation^69^. To adjust for tumor purity and normal contamination, the *–m clonal* option was used with tumor purities from pathologic evaluations. Copy number ratios were then plotted with the *heatmap* function and copy number values were assessed and plotted with *ggplot2*. Gene-level copy number estimates represent the mean copy number call across all probe targets. *CNVkit* copy number ratios showed a near normal distribution and *ERBB2* copy number values demonstrated a strong correlation (pearson R = 0.924, p-value < 0.001) with expression (Appendix A.4: Figure 42).

### Differential gene expression, clustering and outlier gains and losses

Hierarchical clustering was performed using the heatmap.3 function (https://raw.githubusercontent.com/obigriffith/biostar-tutorials/master/Heatmaps/heatmap.3.R) in R on log2normCPM values of the top 10% most variable genes (defined by IQR) with 1 minus Pearson correlations as distance measurements and the “average” agglomeration method. Differential expression between primary and recurrent tumors was analyzed with *limma*. Raw counts were input into the *voom* function and quantile normalized prior to fitting the linear model and performing the empirical Bayes method for differential expression^70,71^. The linear model was fitted with a design that accounts for the paired nature of the cohort (model = ∼Patient+Tissue [primary or recurrence]). Outlier expression gains and losses were determined for each patient by discretely categorizing genes into one of 5 categories. If log2FC values (i.e. recurrence log2normCPM – primary log2normCPM) for a given gene were less than Q1 – (1.5 X IQR) or Q1 – (3 X IQR), using case-specific log2FC values for all genes as the distribution, that gene was deemed an “Outlier Loss” or “Extreme Loss” respectively. If log2FC values calculated were greater than Q3 + (1.5 X IQR) or Q3 + (3 X IQR), it was deemed an “Outlier Gain” or “Extreme Gain” respectively. All other genes with intermediate fold changes were classified as “Stable.” To determine subtype expression of *KLK7, PROM1* and *NDRG1*, normalized microarray expression data along with PAM50 calls was obtained from the Molecular Taxonomy of Breast Cancer International Consortium (METABRIC) through Synapse (https://www.synapse.org/, Synapse ID: syn1688369), following IRB approval for data access from the University of Pittsburgh^72^. Overlap with genes in long-term estrogen deprived, ER-positive breast cancer lines (HCC1428, MCF7, T47D, ZR75.1) was performed by running a separate differential expression analysis (LTED vs. parental lines) on microarray data with *limma*^71,73^. Dysregulated gene overlap was designated if the nominal p-value and FDR-adjusted p-value were both < 0.05 in the local recurrence and LTED differential expression analysis, respectively. Binary dichotomization of METABRIC samples using *NDRG1* expression (>50^th^ percentile, <50^th^ percentile) and log-rank testing were used to assess significant differences in disease-specific survival (DSS) and then Kaplan-Meier curves were plotted with *survminer*^74,75^.

## AUTHOR CONTRIBUTIONS

Study concept and design (NP, KD, SO, AVL); acquisition, analysis, or interpretation of data (all authors); drafting of the manuscript (NP, KD, SO, AVL); critical revision of the manuscript for important intellectual content (all authors); administrative, technical, or material support (WH, JKK, TD, PCL, JUB, CD, AM, BIH)

## ACKNOWLEDGMENTS

This project used the University of Pittsburgh HSCRF Genomics Research Core and Pitt Biospecimen Core, and the UPMC HIllman Cancer Center Tissue and Research Pathology Services supported in part by award P30CA047904. The authors would like to thank the patients who contributed samples to this study and Lori Miller (University of Pittsburgh) for their efforts in collecting tissue.

## Grant Support

Research funding for this project was provided in part by Susan G. Komen Scholar awards (AVL and SO), Breast Cancer Research Foundation (AVL and SO), the Fashion Footwear Association of New York, Magee-Women’s Research Institute and Foundation, Nicole Meloche Foundation, Penguins Foundation, Penguins Alumni Foundation, Mario Lemieux. NP was supported by a training grant from the NIH/NIGMS (2T32GM008424-21), an individual fellowship from the NIH/NCI (5F30CA203095) and University of Pittsburgh School of Medicine Dean’s MSTP Postdoctoral Scholar Award.

## Conflicts of Interest Disclosure

Illumina provided free reagents for study but had no role in study design, data analysis or conclusions. CD reports personal fees from Novartis, Roche, MSD Oncology and Daiichi Sankyo, grants from Myriad Genetics and is a shareholder of Sividon Diagnostics / Myriad. In addition, Dr. Denkert has a patent EP18209672 pending, a patent EP20150702464 pending, and a patent Software (VMscope digital pathology) pending. JUB has received Honoraria from Amgen, AstraZeneca, Pfizer, Novartis, SonoScape, MSD Oncology, Roche and has served a consulting role for Pfizer, Amgen, Novartis, AstraZeneva and Roche.

**SUPPLEMENTARY FIGURES AND LEGENDS ARE PROVIDED IN SEPARATE FILE**

## REFERENCES

1. Early Breast Cancer Trialists’ Collaborative Group (EBCTCG) et al. Relevance of breast cancer hormone receptors and other factors to the efficacy of adjuvant tamoxifen: patient-level meta-analysis of randomised trials. The Lancet 378, 771–784 (2011).

2. Davies, C. et al. Long-term effects of continuing adjuvant tamoxifen to 10 years versus stopping at 5 years after diagnosis of oestrogen receptor-positive breast cancer: ATLAS, a randomised trial. The Lancet 381, 805–816 (2013).

3. Goss, P. E. et al. Extending Aromatase-Inhibitor Adjuvant Therapy to 10 Years. N Engl J Med 375, 209–219 (2016).

4. Jacobson, J. A. et al. Ten-year results of a comparison of conservation with mastectomy in the treatment of stage I and II breast cancer. N Engl J Med 332, 907–911 (1995).

5. Robinson, D. R. et al. Activating ESR1 mutations in hormone-resistant metastatic breast cancer. Nat Genet 45, 1446–1451 (2013).

6. Toy, W. et al. ESR1 ligand-binding domain mutations in hormone-resistant breast cancer. Nat Genet 45, 1439–1445 (2013).

7. Jeselsohn, R., Buchwalter, G., De Angelis, C., Brown, M. & Schiff, R. ESR1 mutations—a mechanism for acquired endocrine resistance in breast cancer. Nat Rev Clin Oncol 12, 573–583 (2015).

8. Kuukasjärvi, T., Kononen, J., Helin, H., Holli, K. & Isola, J. Loss of estrogen receptor in recurrent breast cancer is associated with poor response to endocrine therapy. J Clin Oncol 14, 2584–2589 (1996).

9. Oh, A. S. et al. Hyperactivation of MAPK induces loss of ERalpha expression in breast cancer cells. Mol Endocrinol 15, 1344–1359 (2001).

10. Creighton, C. J. et al. Activation of mitogen-activated protein kinase in estrogen receptor alpha-positive breast cancer cells in vitro induces an in vivo molecular phenotype of estrogen receptor alpha-negative human breast tumors. Cancer Res 66, 3903–3911 (2006).

11. Shou, J. et al. Mechanisms of Tamoxifen Resistance: Increased Estrogen Receptor-HER2/neu Cross-Talk in ER/HER2-Positive Breast Cancer. JNCI Journal of the National Cancer Institute 96, 926–935 (2004).

12. Turner, N. et al. FGFR1 amplification drives endocrine therapy resistance and is a therapeutic target in breast cancer. Cancer Res 70, 2085–2094 (2010).

13. Basudan, A. et al. Frequent ESR1 and CDK Pathway Copy-Number Alterations in Metastatic Breast Cancer. Mol Cancer Res 17, 457–468 (2019).

14. Hartmaier, R. J. et al. Recurrent hyperactive ESR1 fusion proteins in endocrine therapy-resistant breast cancer. Ann Oncol 29, 872–880 (2018).

15. Gundem, G. et al. The evolutionary history of lethal metastatic prostate cancer. Nature 520, 353–357 (2015).

16. Hugo, W. et al. Non-genomic and Immune Evolution of Melanoma Acquiring MAPKi Resistance. Cell 162, 1271–1285 (2015).

17. Nik-Zainal, S. et al. The life history of 21 breast cancers. Cell 149, 994–1007 (2012).

18. Hanker, A. B. et al. An Acquired HER2T798I Gatekeeper Mutation Induces Resistance to Neratinib in a Patient with HER2 Mutant-Driven Breast Cancer. Cancer Discov 7, 575–585 (2017).

19. Miller, W. R. et al. Changes in breast cancer transcriptional profiles after treatment with the aromatase inhibitor, letrozole. Pharmacogenet Genomics 17, 813–826 (2007).

20. Mackay, A. et al. Molecular response to aromatase inhibitor treatment in primary breast cancer. Breast Cancer Res 9, R37 (2007).

21. Gutierrez, M. C. et al. Molecular changes in tamoxifen-resistant breast cancer: relationship between estrogen receptor, HER-2, and p38 mitogen-activated protein kinase. J. Clin. Oncol. 23, 2469–2476 (2005).

22. Varešlija, D. et al. Adaptation to AI Therapy in Breast Cancer Can Induce Dynamic Alterations in ER Activity Resulting in Estrogen-Independent Metastatic Tumors. Clin Cancer Res 22, 2765–2777 (2016).

23. Miller, C. A. et al. Aromatase inhibition remodels the clonal architecture of estrogen-receptor-positive breast cancers. Nat Commun 7, 12498 (2016).

24. Razavi, P. et al. The Genomic Landscape of Endocrine-Resistant Advanced Breast Cancers. Cancer Cell 34, 427–438.e6 (2018).

25. Nayar, U. et al. Acquired HER2 mutations in ER+ metastatic breast cancer confer resistance to estrogen receptor-directed therapies. Nat Genet 51, 207–216 (2019).

26. Pearson, A. et al. Inactivating NF1 mutations are enriched in advanced breast cancer and contribute to endocrine therapy resistance. Clin Cancer Res 26, 608–622 (2020).

27. Zheng, Z.-Y. et al. Neurofibromin Is an Estrogen Receptor-α Transcriptional Co-repressor in Breast Cancer. Cancer Cell 37, 387–402.e7 (2020).

28. Priedigkeit, N. et al. Exome-capture RNA sequencing of decade-old breast cancers and matched decalcified bone metastases. JCI Insight 2, (2017).

29. Forbes, S. A. et al. COSMIC: exploring the world’s knowledge of somatic mutations in human cancer. Nucleic Acids Res 43, D805–11 (2015).

30. Du, T. et al. Key regulators of lipid metabolism drive endocrine resistance in invasive lobular breast cancer. Breast Cancer Res 20, 106 (2018).

31. Wang, C., Christin, J. R., Oktay, M. H. & Guo, W. Lineage-Biased Stem Cells Maintain Estrogen-Receptor-Positive and -Negative Mouse Mammary Luminal Lineages. Cell Rep 18, 2825–2835 (2017).

32. Gao, R. et al. Punctuated copy number evolution and clonal stasis in triple-negative breast cancer. Nat Genet 48, 1119–1130 (2016).

33. Wahner-Roedler, D. L., Reynolds, C. A. & Boughey, J. C. Collision tumors with synchronous presentation of breast carcinoma and lymphoproliferative disorders in the axillary nodes of patients with newly diagnosed breast cancer: a case series. Clin Breast Cancer 11, 61–66 (2011).

34. Liu, L., Kimball, S., Liu, H., Holowatyj, A. & Yang, Z.-Q. Genetic alterations of histone lysine methyltransferases and their significance in breast cancer. Oncotarget 6, 2466–2482 (2015).

35. Pereira, B. et al. The somatic mutation profiles of 2,433 breast cancers refines their genomic and transcriptomic landscapes. Nat Commun 7, 11479 (2016).

36. Manso, L. et al. Analysis of Paired Primary-Metastatic Hormone-Receptor Positive Breast Tumors (HRPBC) Uncovers Potential Novel Drivers of Hormonal Resistance. PLoS ONE 11, e0155840 (2016).

37. Jones, S. et al. Frequent mutations of chromatin remodeling gene ARID1A in ovarian clear cell carcinoma. Science 330, 228–231 (2010).

38. Guan, B., Wang, T.-L. & Shih, I.-M. ARID1A, a factor that promotes formation of SWI/SNF-mediated chromatin remodeling, is a tumor suppressor in gynecologic cancers. Cancer Res 71, 6718–6727 (2011).

39. Zhang, X. et al. Frequent low expression of chromatin remodeling gene ARID1A in breast cancer and its clinical significance. Cancer Epidemiol 36, 288–293 (2012).

40. Xu, G. et al. ARID1A determines luminal identity and therapeutic response in estrogen-receptor-positive breast cancer. Nat Genet 52, 198–207 (2020).

41. Saito, Y. et al. Landscape and function of multiple mutations within individual oncogenes. Nature (2020). doi: 10.1038/s41586-020-2175-2

42. Zundelevich, A. et al. ESR1 mutations are frequent in newly diagnosed metastatic and loco-regional recurrence of endocrine-treated breast cancer and carry worse prognosis. Breast Cancer Res 22, 16 (2020).

43. Jeselsohn, R. et al. Allele-Specific Chromatin Recruitment and Therapeutic Vulnerabilities of ESR1 Activating Mutations. Cancer Cell 33, 173–186.e5 (2018).

44. Drilon, A. et al. Efficacy of Larotrectinib in TRK Fusion-Positive Cancers in Adults and Children. N Engl J Med 378, 731–739 (2018).

45. Veeck, J. et al. Promoter hypermethylation of the SFRP2 gene is a high-frequent alteration and tumor-specific epigenetic marker in human breast cancer. Mol Cancer 7, 83 (2008).

46. Loh, Y. N. et al. The Wnt signalling pathway is upregulated in an in vitro model of acquired tamoxifen resistant breast cancer. BMC Cancer 13, 174 (2013).

47. Sikora, M. J. et al. WNT4 mediates estrogen receptor signaling and endocrine resistance in invasive lobular carcinoma cell lines. Breast Cancer Res 18, 92 (2016).

48. Touat, M., Ileana, E., Postel-Vinay, S., André, F. & Soria, J.-C. Targeting FGFR signaling in cancer. Clin Cancer Res 21, 2684–2694 (2015).

49. Levine, K. M. et al. FGFR4 overexpression and hotspot mutations in metastatic ER+ breast cancer are enriched in the lobular subtype. NPJ Breast Cancer 5, 19 (2019).

50. Subramanian, A. et al. Gene set enrichment analysis: a knowledge-based approach for interpreting genome-wide expression profiles. Proc Natl Acad Sci U S A 102, 15545–15550 (2005).

51. Sansone, P. et al. Self-renewal of CD133(hi) cells by IL6/Notch3 signalling regulates endocrine resistance in metastatic breast cancer. Nat Commun 7, 10442 (2016).

52. Sansone, P. et al. Evolution of Cancer Stem-like Cells in Endocrine-Resistant Metastatic Breast Cancers Is Mediated by Stromal Microvesicles. Cancer Res 77, 1927–1941 (2017).

53. Guan, R. J. et al. Drg-1 as a differentiation-related, putative metastatic suppressor gene in human colon cancer. Cancer Res 60, 749–755 (2000).

54. Patro, R., Duggal, G., Love, M. I., Irizarry, R. A. & Kingsford, C. Salmon provides fast and bias-aware quantification of transcript expression. Nat Methods 14, 417–419 (2017).

55. Soneson, C., Love, M. I. & Robinson, M. D. Differential analyses for RNA-seq: transcript-level estimates improve gene-level inferences. [version 2; peer review: 2 approved]. F1000Res 4, 1521 (2016).

56. Robinson, M. D., McCarthy, D. J. & Smyth, G. K. edgeR: a Bioconductor package for differential expression analysis of digital gene expression data. Bioinformatics 26, 139–140 (2010).

57. Robinson, M. D. & Oshlack, A. A scaling normalization method for differential expression analysis of RNA-seq data. Genome Biol 11, R25 (2010).

58. Li, H. & Durbin, R. Fast and accurate short read alignment with Burrows-Wheeler transform. Bioinformatics (Oxford, England) 25, 1754–1760 (2009).

59. Li, H. et al. The Sequence Alignment/Map format and SAMtools. Bioinformatics (Oxford, England) 25, 2078–2079 (2009).

60. McKenna, A. et al. The Genome Analysis Toolkit: a MapReduce framework for analyzing next-generation DNA sequencing data. Genome Res 20, 1297–1303 (2010).

61. Koboldt, D. C. et al. VarScan 2: somatic mutation and copy number alteration discovery in cancer by exome sequencing. Genome Res 22, 568–576 (2012).

62. Obenchain, V. et al. VariantAnnotation: a Bioconductor package for exploration and annotation of genetic variants. Bioinformatics 30, 2076–2078 (2014).

63. Wong, S. Q. et al. Sequence artefacts in a prospective series of formalin-fixed tumours tested for mutations in hotspot regions by massively parallel sequencing. BMC Med Genomics 7, 23 (2014).

64. Wang, K., Li, M. & Hakonarson, H. ANNOVAR: functional annotation of genetic variants from high-throughput sequencing data. Nucleic Acids Res 38, e164 (2010).

65. Lek, M. et al. Analysis of protein-coding genetic variation in 60,706 humans. Nature 536, 285–291 (2016).

66. Gu, Z., Eils, R. & Schlesner, M. Complex heatmaps reveal patterns and correlations in multidimensional genomic data. Bioinformatics (Oxford, England) btw313 (2016).

67. Robinson, J. T. et al. Integrative genomics viewer. Nat Biotechnol 29, 24–26 (2011).

68. Talevich, E., Shain, A. H., Botton, T. & Bastian, B. C. CNVkit: Genome-Wide Copy Number Detection and Visualization from Targeted DNA Sequencing. PLoS Comput Biol 12, e1004873 (2016).

69. Olshen, A. B., Venkatraman, E. S., Lucito, R. & Wigler, M. Circular binary segmentation for the analysis of array-based DNA copy number data. Biostatistics 5, 557–572 (2004).

70. Law, C. W., Chen, Y., Shi, W. & Smyth, G. K. voom: Precision weights unlock linear model analysis tools for RNA-seq read counts. Genome Biol 15, R29 (2014).

71. Ritchie, M. E. et al. limma powers differential expression analyses for RNA-sequencing and microarray studies. Nucleic Acids Res 43, e47 (2015).

72. Curtis, C. et al. The genomic and transcriptomic architecture of 2,000 breast tumours reveals novel subgroups. Nature 486, 346–352 (2012).

73. Simigdala, N. et al. Cholesterol biosynthesis pathway as a novel mechanism of resistance to estrogen deprivation in estrogen receptor-positive breast cancer. Breast Cancer Res 18, 58 (2016).

74. Bland, J. M. & Altman, D. G. The logrank test. BMJ 328, 1073 (2004).

75. Kassambara, A. & Kosinski, M. survminer: Drawing Survival Curves using “ggplot2.” (CRAN, 2016).

