## Supplemental Figures for "Acquired mutations and transcriptional remodeling in long-term estrogen-deprived locoregional breast cancer recurrences"

***Supplementary Figure 1.*** DNA-seq target interval coverages

***Supplementary Figure 2.*** Copy number call distribution

***Supplementary Figure 3***. tumorMatch: Proportion of shared variants (POSV) between samples in patient-matched cohort

***Supplementary Figure 4.*** Local recurrence CNA Correlation Matrix

***Supplementary Figure 5.*** Allele frequency of SNVs called from RNA-seq vs DNA-seq

***Supplementary Figure 6.*** Overlap of differentially expressed genes between local recurrences and ER+ LTED lines

***Supplementary Figure 7.*** *KLK7* and *PROM1* basal breast carcinoma expression

***Supplementary Figure 8.*** NDRG1 expression in PAM50 subtypes and survival influence in ER-positive breast cancer

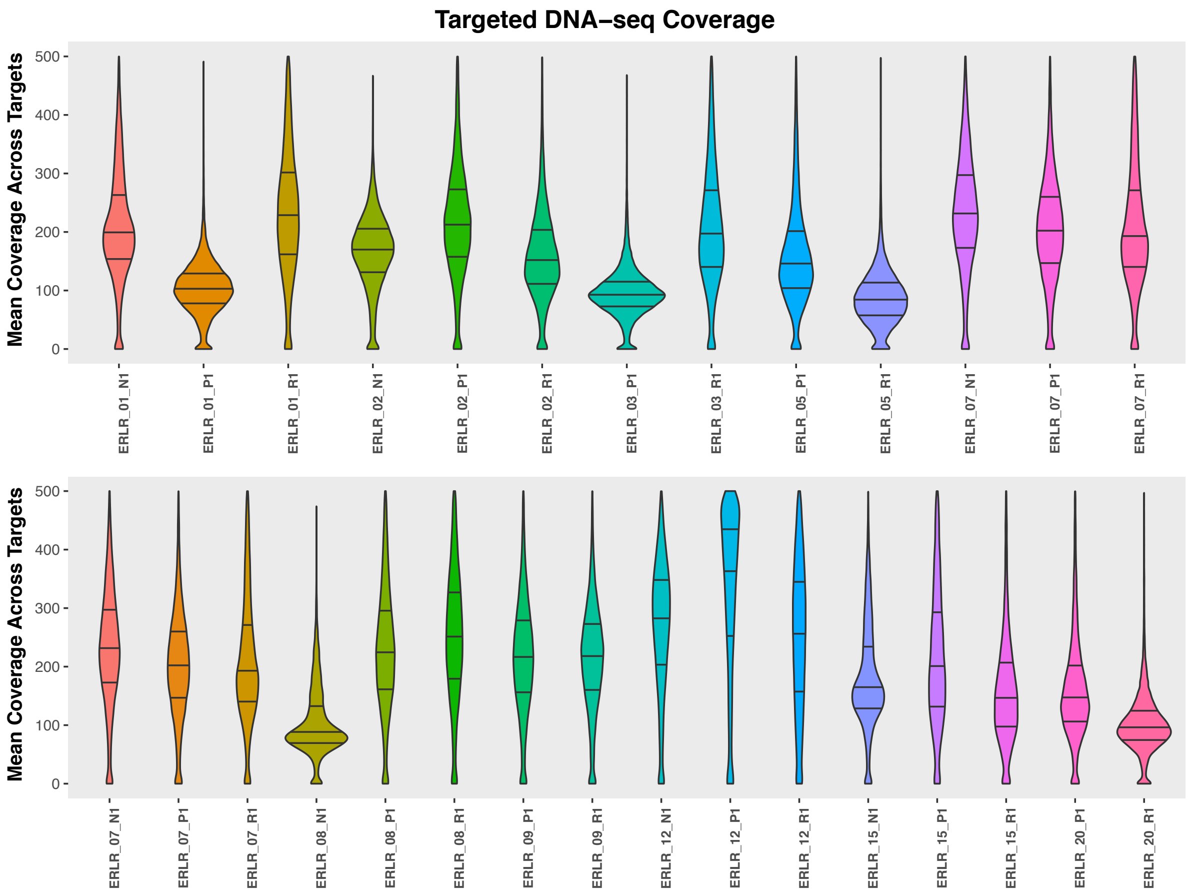


***Supplementary Figure 1:*** ***DNA-seq target interval coverages.*** Violin plots showing the distribution of mean DNA-sequencing coverage across all targeted intervals. 25th, 50th and 75th percentiles are indicated with horizontal black lines. To better visualize distributions, y-axis limit was set at 500.

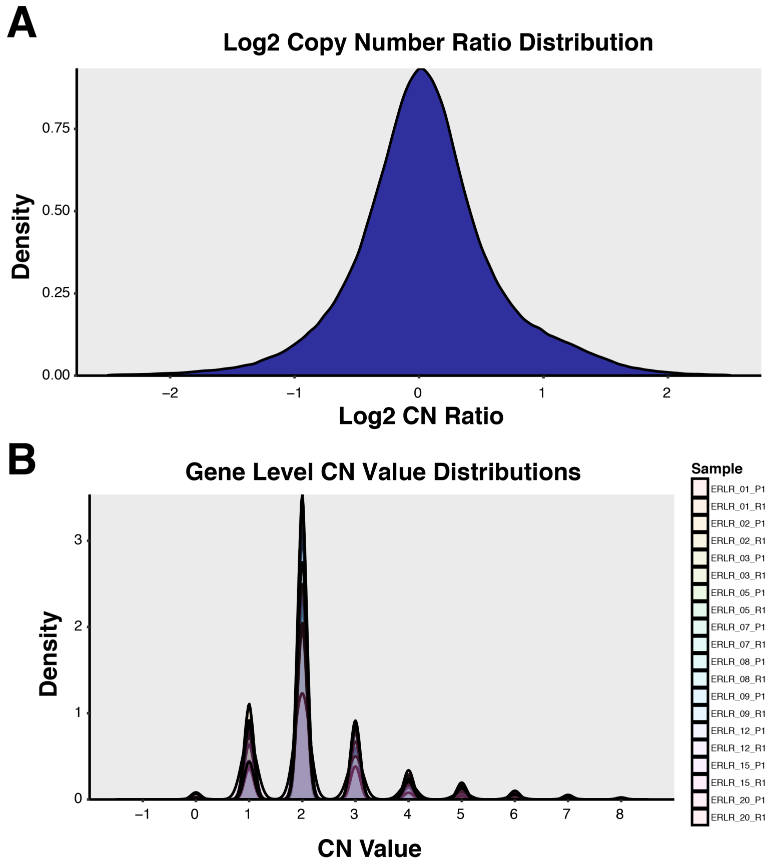


***Supplementary Figure 2:* Copy number call distribution.** **(A)** Log2 copy number ratio distribution, derived from *CNVkit*, for all samples in cohort. **(B)** Distribution of discrete, gene-level copy number calls with gene-level values representing the mean of discrete calls across all probed regions covering the gene.

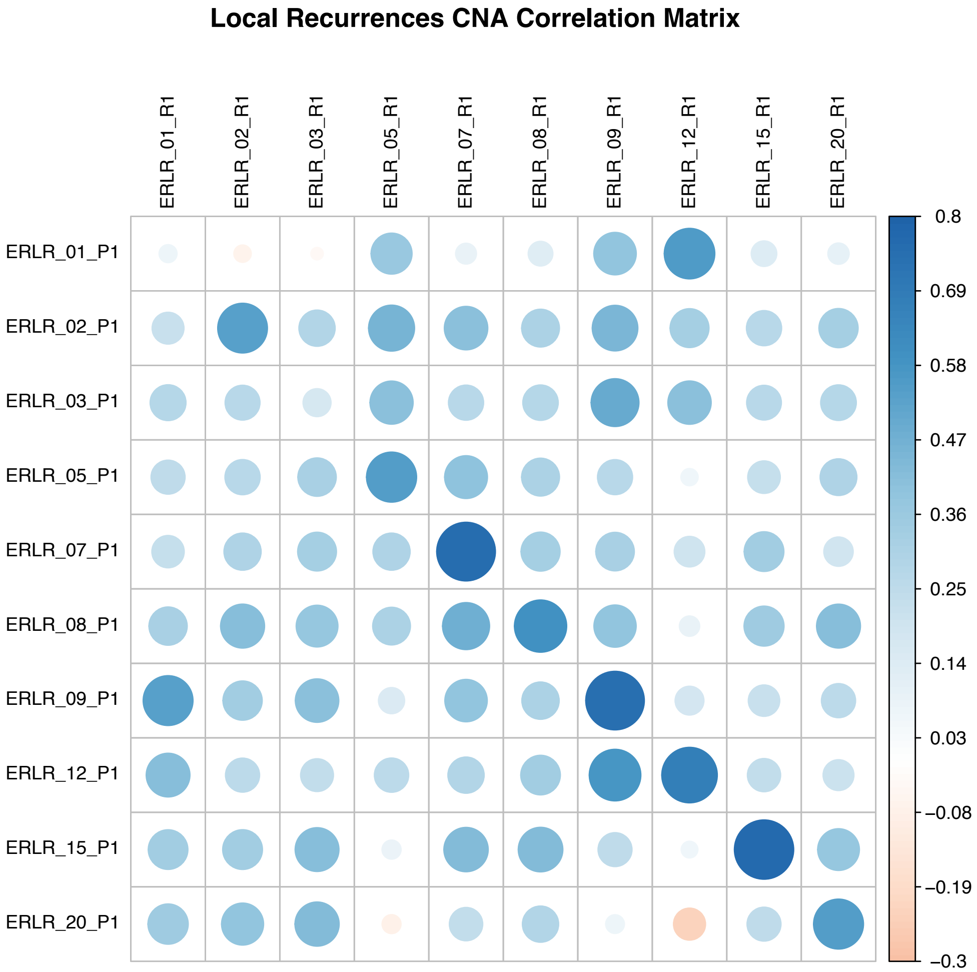


***<u>Supplementary Figure 3:</u> CNA Correlation Matrix.*** Correlation matrix of pearson R values from log2CN values between all samples.

### ***tumorMatch*** **in local recurrence cohort**

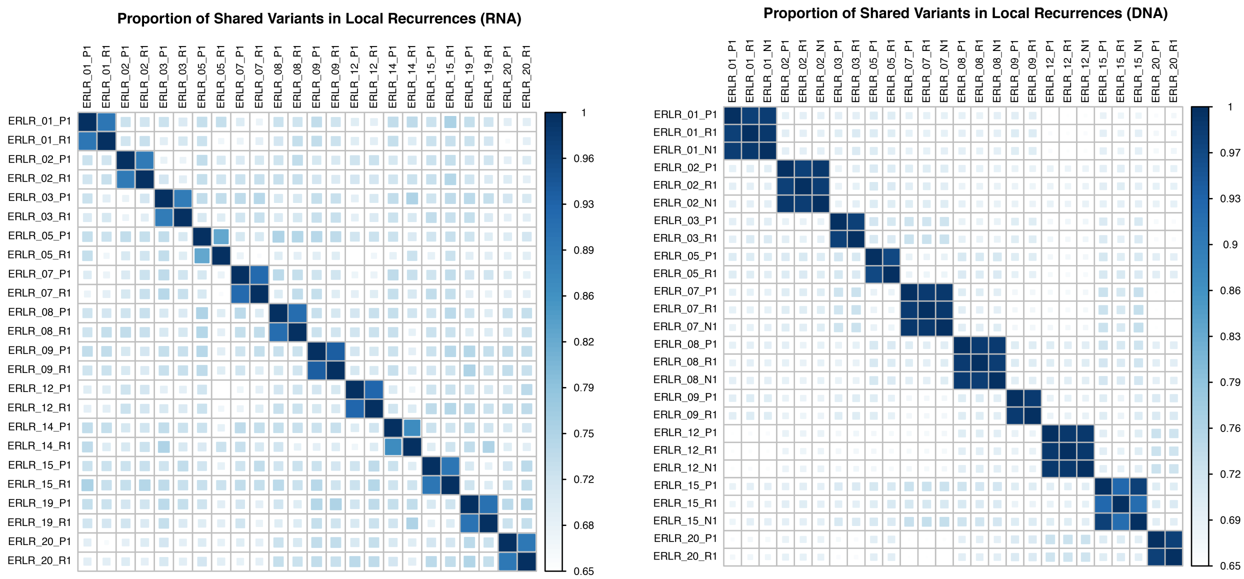


***<u>Supplementary Figure 4:</u> tumorMatch proportion of shared variants (POSV) between samples in patient-matched cohort.*** *tumorMatch* plots for both RNA- (left) and DNA-sequencing, showing all paired specimens, including trios that include normal, are patient-matched.

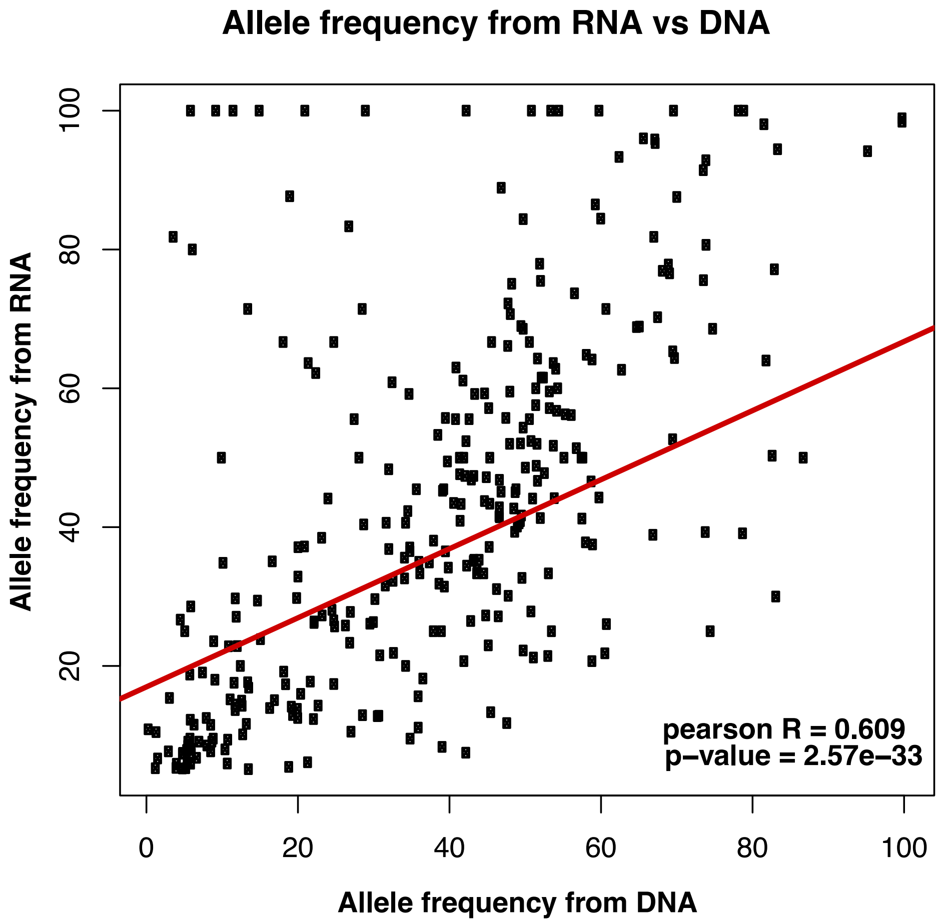


***Supplementary Figure 5:*** **Allele frequency of SNVs called from RNA-seq vs DNA-seq.**

Allele frequency calls of 633 total somatic, nonsynonymous nucleotide variants from RNA-seq and DNA-seq (pearson R = 0.609).

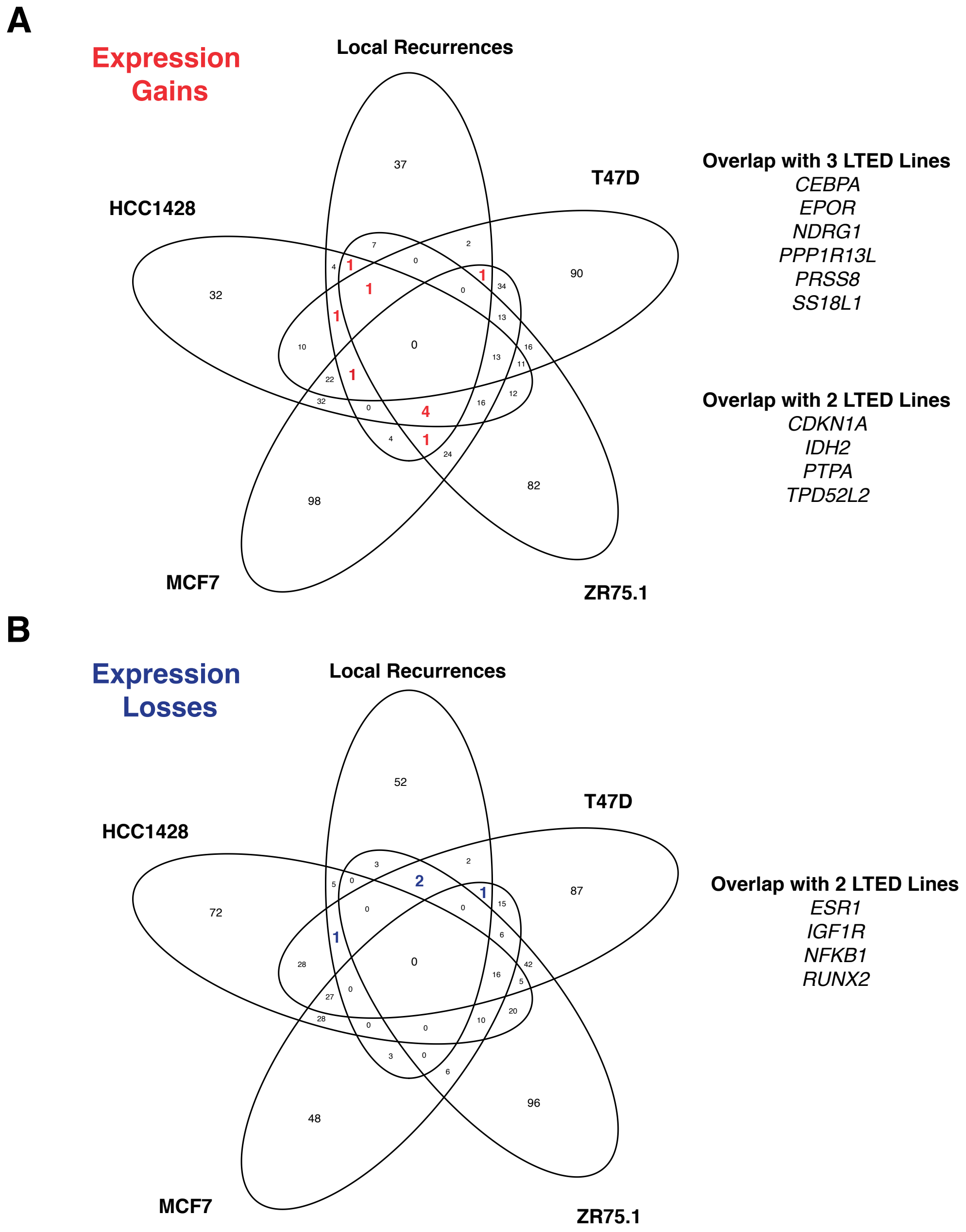


***Supplementary Figure 6:*** **Overlap of differentially expressed genes between local recurrences and ER+ LTED lines. (A)** Genes significantly upregulated in both local recurrences vs. primaries and LTED vs. parental lines. **(B)** Genes significantly downregulated in both local recurrences vs. primaries and LTED vs. parental lines.

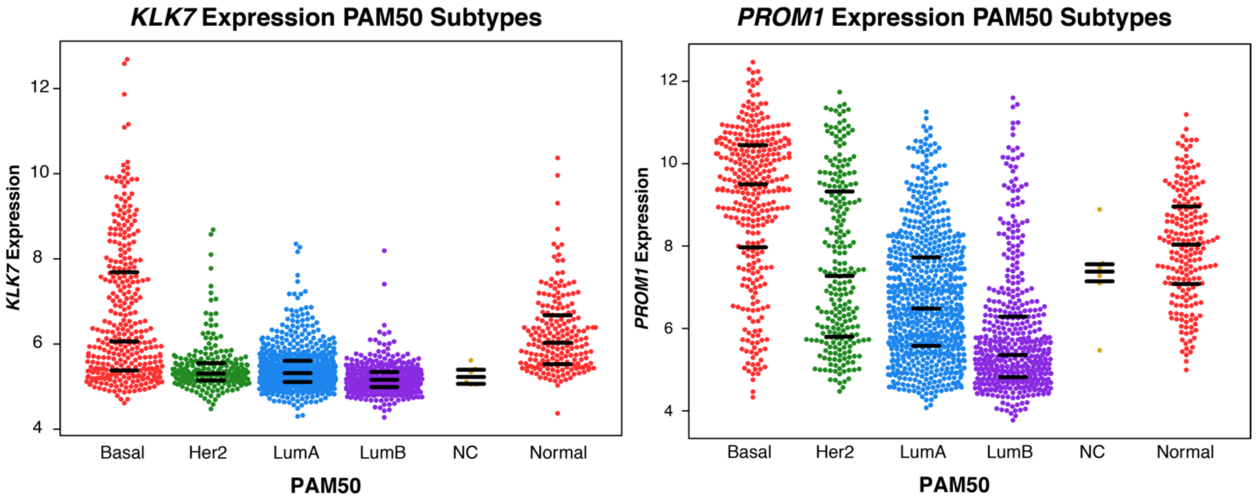


***Supplementary Figure 7:*** ***KLK7*** **and** ***PROM1*** **basal breast carcinoma expression.**

Normalized microarray expression values (METABRIC) of *KLK7* and *PROM1*, segregated by PAM50 subtypes. Horizontal black bars indicate 25$^{th}$, 50$^{th}$ and 75$^{th}$ percentile values.

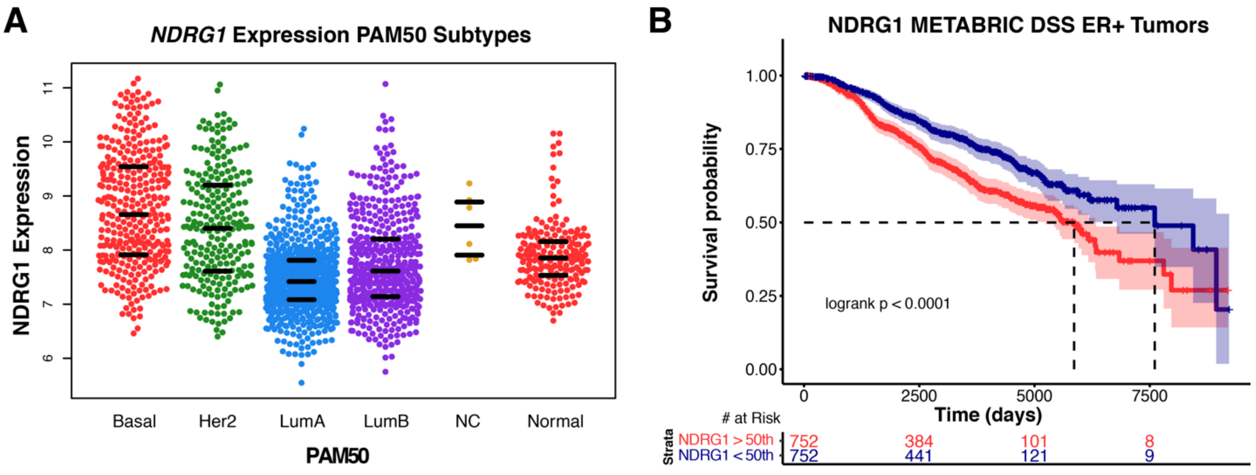


***Supplementary Figure 8:*** **NDRG1 expression in PAM50 subtypes and survival influence in ER-positive breast cancer. (A)** *NDRG1* expression in PAM50 subtypes. **(B)** Disease-specific survival in METABRIC of patients with ER-positive primary tumors that express *NDRG1* highly (>50$^{th}$ percentile, red) or lowly (<50$^{th}$ percentile, blue).
